# Multilayered N-glycoproteomics reveals impaired N-glycosylation promoting Alzheimer’s disease

**DOI:** 10.1101/615989

**Authors:** Pan Fang, Juan-Juan Xie, Shao-Ming Sang, Lei Zhang, Ming-Qi Liu, Lu-Jie Yang, Yi-Teng Xu, Guo-Quan Yan, Jun Yao, Xing Gao, Wen-Jing Qian, Zhong-Feng Wang, Yang Zhang, Peng-Yuan Yang, Hua-Li Shen

**Author notes:** These authors contributed equally to this work (Pan Fang, Juan-Juan Xie). Corresponding authors: Y Zhang, P-Y Yang and H-L Shen, Address: Institutes of Biomedical Sciences of Shanghai Medical School, Fudan University, 131 Dong’an Road, Shanghai 200032, China., (YZ); (PYY); (HLS).

## Abstract

Alzheimer’s disease (AD) is one of the most common neurodegenerative diseases that currently lacks clear pathogenesis and effective treatment. Protein glycosylation is ubiquitous in brain tissue and site-specific analysis of N-glycoproteome, which is technically challenging, can advance our understanding of the glycoproteins’ role in AD. In this study, we profiled the multilayered variations in proteins, N-glycosites, N-glycans, and in particular site-specific N-glycopeptides in the APP/PS1 and wild type mouse brain through combining pGlyco 2.0 strategy with other quantitative N-glycoproteomic strategies. The comprehensive brain N-glycoproteome landscape was constructed, and rich details of the heterogeneous site-specific protein N-glycosylations were exhibited. Quantitative analyses explored generally downregulated N-glycosylation involving proteins such as glutamate receptors, as well as fucosylated and oligo-mannose type glycans in APP/PS1 mice versus wild type mice. Moreover, our preliminary functional study revealed that N-glycosylation was crucial for the membrane localization of NCAM1 and for maintaining the excitability and viability of neuron cells. Our work offered a panoramic view of the N-glycoproteomes in Alzheimer’s disease and revealed that generally impaired N-glycosylation promotes Alzheimer’s disease progression.

## INTRODUCTION

Alzheimer’s disease is one of the most common dementias that currently affects more than 47 million people worldwide(Haapasalo *et al*, 2015). Unfortunately, no effective treatments are available to prevent, halt, or reverse Alzheimer’s disease, and its pathogenic mechanisms remain unclear(De Strooper and Karran, 2016; Huang and Mucke, 2012; Selkoe and Hardy, 2016).

Glycosylation is one of the most universal and essential modifications. It determines protein folding, trafficking, stability, and regulates many cellular activities(Xiao *et al*, 2019). Our previous work has explored that protein glycosylation is ubiquitous in mouse brain and the involved proteins are vital for brain functions, including memory and learning(Fang *et al*, 2016). Some correlations were found between protein N-glycosylation and Alzheimer’s disease(Schedin-Weiss *et al*, 2014a). Abnormal glycan profilings were observed in Alzheimer’s disease patients(Chen *et al*, 2010; Frenkel-Pinter *et al*, 2017; Gizaw *et al*, 2016; Palmigiano *et al*, 2016). Furthermore, Many Alzheimer’s disease-related proteins, including APP, tau, BACE1, nicastrin and so on, were found to be glycosylated, and their glycosylation patterns changed in Alzheimer’s disease pathogenesis(Kizuka *et al*, 2015; Kizuka *et al*, 2017; Schedin-Weiss *et al*, 2014b). Thus, a systematic study on the global changes of the N-glycosylated proteins, especially their site-specific variations will link these scattered findings and contribute to expand our understanding of glycosylation in Alzheimer’s disease.

The direct identification and quantification of N-glycopeptides are indispensable for a systematic N-glycoproteomic study. Recently, N-glycoproteomics techniques on high-sensitive glycopeptides enrichment(Fang *et al*, 2016) and high-throughput site-specific N-glycopeptides identification have experienced great progresses(Ruhaak *et al*, 2018; Stadlmann *et al*, 2017; Sun *et al*, 2016; Zeng *et al*, 2016). In particular, we developed a large scale and precision identification method for site-specific N-glycopeptides named pGlyco 2.0(Liu *et al*, 2017).

In this work, through applying pGlyco 2.0 for the intact N-glycopeptides, iTRAQ labeling quantification for the whole proteome and the deglycopeptides, as well as lectin microarray for glycan epitopes, a multilayered N-glycoproteome landscape was constructed in the APP/PS1 double-transgenic mouse versus wild type mouse brain. General and site-specific variations of the protein N-glycosylations in the APP/PS1 mice and wild type mice were revealed. Furthermore, influences of the varied glycosylation on NCAM1 and hippocampal neurons were explored. This work provides a panoramic view of protein N-glycosylation changes in APP/PS1 mice versus wild type mice and could present novel insights into the molecular mechanisms in the Alzheimer’s disease pathogenesis.

## RESULTS

### 1. Construction of the quantitative proteome, N-glycoproteome, and glycome landscape in the APP/PS1 mouse and wild type mouse brain

Multiple technologies were integrated to quantify the proteome, N-glycoproteome, and glycome in the APP/PS1 mouse and wild type mouse brain (**Fig. 1A**). Quantitative proteomic analysis using iTRAQ labeling and 2D LC-MS/MS led to the quantification of 5,046 proteins in APP/PS1 mice and wild type mice in two biological and two technical replicates. In N-glycosite quantification, the iTRAQ labeled glycopeptides were enriched by ZIC-HILIC followed by the detachment of N-glycans. A total of 804 N-glycosites on 375 proteins were quantified in two biological replicates and three technical replicates for wild type mice group, and one biological replicate and three technical replicates for APP/PS1 mice. Meanwhile, intact N-glycopeptides after ZIC-HILIC enrichment were directly identified with pGlyco 2.0 strategy and quantified using label-free method, and 3,524 intact glyco-peptides were identified from the two group of samples in three biological replicates. The proteins extracted from brain tissues were also subjected to lectin microarrays containing 91 lectins with distinct glycan binding specificities and 83 lectins were positively detected (**Fig. 1A and E**). Through combing above data, the N-glycoproteome landscape of mouse brain consisting of 5,322 proteins (including 722 glycoproteins), 1493 N-glycosites, 3,524 glyco-peptides, and 83 lectin recognized glycan patterns was constructed (**Fig. 1B**). Of the identified proteins, 276 were exclusively contributed by the N-glycoproteome data (**Fig. 1C**). Among the total 1,493 N-glycosites, 423 and 689 N-glycosites were only identified by “N-glycosites” and “site-specific N-glycopeptides” strategies, respectively, while 381 glycosites were co-identified by the two strategies (**Fig. 1D**). The multi-omic data provided a panoramic view of the proteomes, N-glycoproteomes and glycomes in the APP/PS1 mouse and wild type mouse brain.

**Figure 1.**
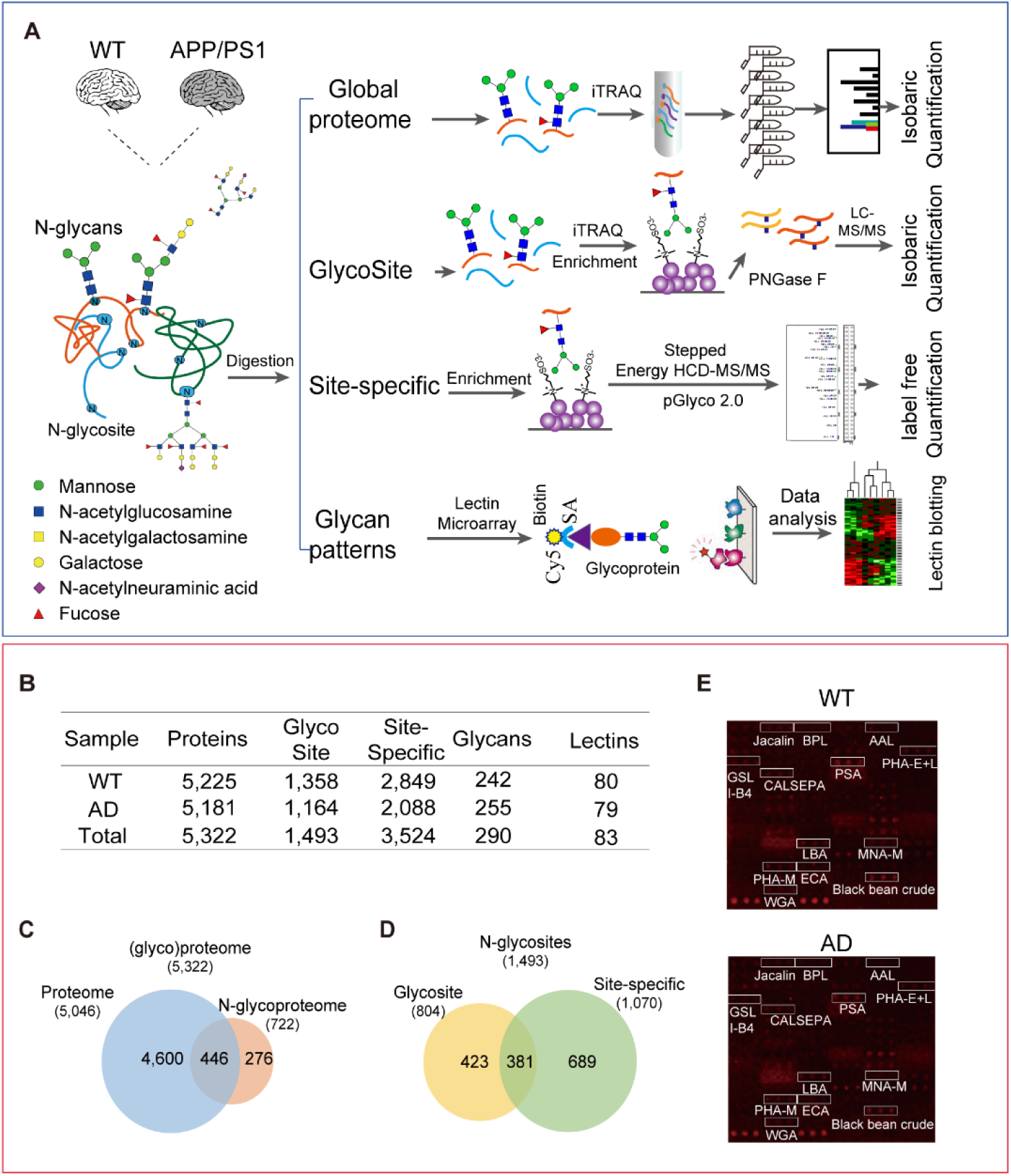
Graphic abstract of the multilayered N-glycoproteomics in the APP/PS1 mouse and wild type mouse brain. **(A)** Quantitative analysis of global proteome, N-glycosites, site-specific N-glycopeptides, and glycan patterns were applied. The acquired multi-omics data were integrated for analyzing the variations in the N-glycoproteome and glycan profiling in APP/PS1 mice (AD) versus wild type mice (WT). N-glycan structures: Green circle, mannose; blue square, N-acetylglucosamine; yellow square, N-acetylgalactosamine; yellow circle, galactose; purple diamond, N-acetylneuraminic acid; red triangle, fucose. **(B)** An overall identification of the proteome, N-glycoproteome, and glycome in AD and WT mouse brain. **(C)** Protein identifications in the brain proteome and N-glycoproteome. **(D)** N-glycosites identifications from the “N-glycosite” and “Site-specific N-glycopeptide” experiments. **(E)** The fluorescent images of Cy5-labeled brain proteins from the WT and AD mouse bound to lectin microarrays. Lectins marked in a white frame indicate the significantly differential signaling intensities between AD and WT.

### 2. Complex and heterogeneous N-glycoproteome landscape of the APP/PS1 mouse and wild type mouse brain

The micro-heterogeneity of protein N-glycosylation was exhibited through analyzing the distribution of glycosites and site-specific glycopeptides on the glycoproteins. About 58.4% of N-glycoproteins have one N-glycosite, while the 7.6% most severely N-glycosylated proteins have more than 5 sites **(Fig. 2A)**. The average number of N-glycosites per glycoprotein is 2. The distribution of N-glycoforms on each site was shown in **Fig. 2B**. The average number of glycoforms per site is 3.3, while there are 35 N-glycosites had more than 14 N-glycans on each of them. Proteins having the most glycosites and the most heterogeneous sites were listed in the Figure (**Fig. 2A and B**). Combining both information, the most heterogeneous glycoproteins in mouse brain were listed in **Figure EV1A**. AT1B2, THY1, EAA2, and NPTN exhibited the severest micro-heterogeneity, with more than100 N-glycans on each protein (**Figure EV1A**). The prolow-density lipoprotein receptor-related protein 1 (LRP1_MOUSE) had 30 N-glycosites (**Fig. 2A**) and was one of the most heavily N-glycosylated protein in our dataset (**Figure EV1A**). The overall distribution of the N-glycopeptides with different lengths (the number of monosaccharides) of N-glycans between the APP/PS1 mice and wild type mice were comparable. The most frequent N-glycans consisted of 7-13 monosaccharides (**Fig. 2C and Figure EV1C)**, while a lot of very long and complex **N-**glycans (14-21 monosaccharides) existed in the APP/PS1 mice and wild type mice **(Fig. 2C and Figure EV1D)**. High-quality MS/MS spectra of the N-glycopeptides with long N-glycans were exhibited (**Fig. 2D and Figure EV1F**). Our method showed strong ability in identifying the N-glycopeptides with highly complex N-glycan structures.

**Figure 2.**
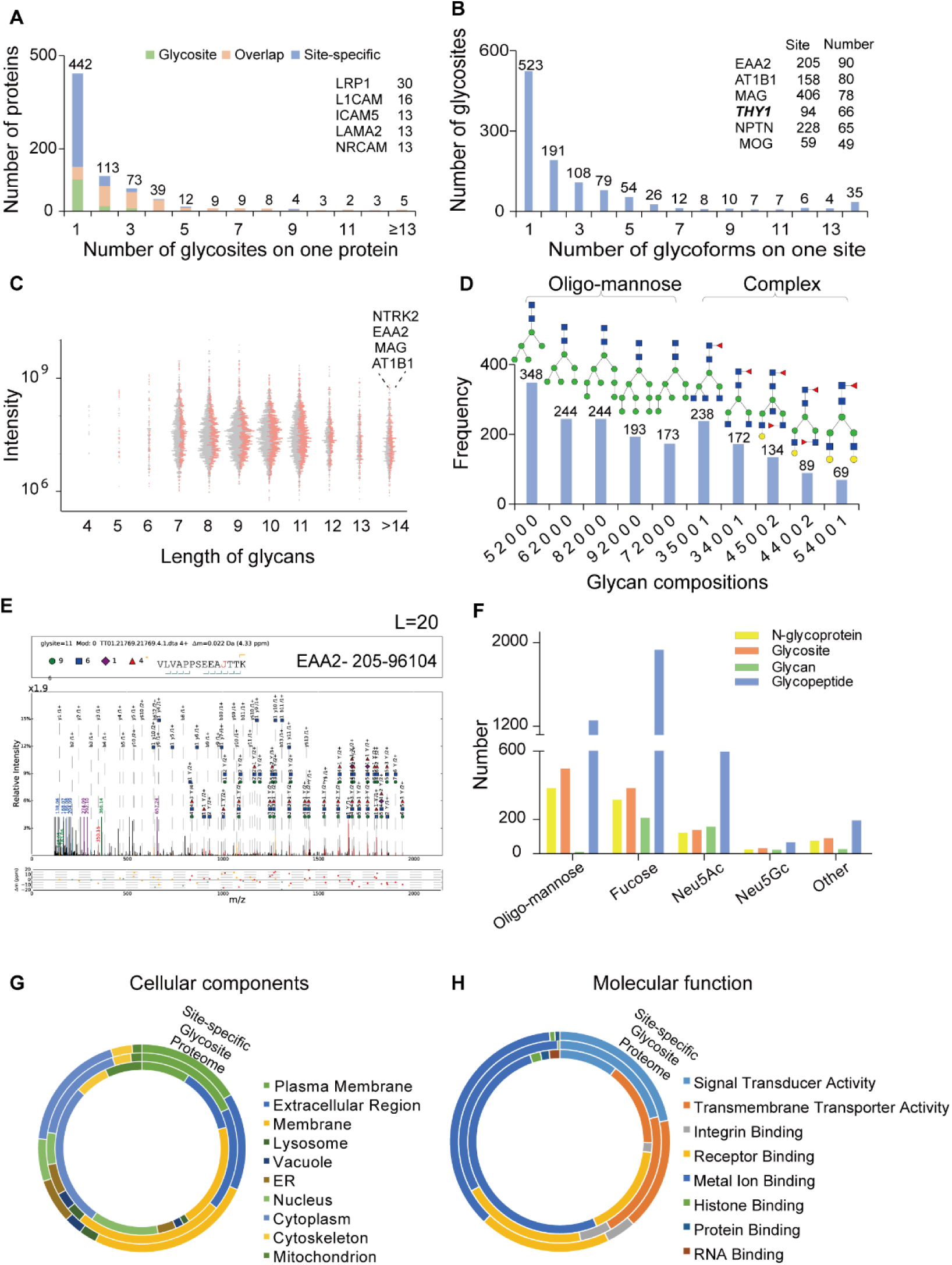
Comprehensive characterization of protein N-glycosylations in the APP/PS1 mouse and wild type mouse brain. **(A)** Frequency distribution of proteins with different numbers of N-glycosites. **(B)** Frequency distribution of N-glycosites with different number of N-glycans. **(C)** Beam worm plot displays the intensities of N-glycopeptides with different lengths of N-glycans in AD and WT mouse brain, orange dots represent the N-glycopeptides from AD and gray dots represent those from WT. **(D)** Ten N-glycans of the highest frequencies on site-specific N-glycopeptides. The composition of an N-glycan on x axis is represented by the number of Hex, HexNAc, NeuAc, NeuGc, and Fuc. The inferred structures of N-glycans were depicted on top of the bars. **(E)** Example MS/MS spectrum of the N-glycopeptide “VLVAPPSEEAJ(96104)TTK” of EAA2_MOUSE proteins with the highly complex and branched N-glycans on its 205 site. “J” indicated glycosite N. **(F)** Frequency distribution of N-glycoprotein, N-glycosite, N-glycan and site-specific glycopeptides with different N-glycan subcategories. **(G)** The cellular localization distribution of the proteins and N-glycoproteins from proteome, N-glycosite, site-specific N-glycopeptide data. **(H)** The molecular function distribution of the proteins and N-glycoproteins from proteome, N-glycosite, site-specific N-glycopeptide data.

A close inspection of N-glycan subcategories showed that N-glycoproteins, N-glycosites and site-specific N-glycopeptides in mouse brain prefer oligo-mannose or fucosylated N-glycans (**Fig. 2E**). In addition, ten most popular N-glycans were attached to approximately 55% (1,945/3,524) of the site-specific N-glycopeptides, suggesting the preferences of these N-glycans on mouse brain proteins (**Fig. 2F**). It is worth noting that these highly frequent N-glycans were all belonged to oligo-mannose or complex type fucosylated N-glycans. Overall, oligo-mannose and fucosylated N-glycans were highly expressed in the APP/PS1 mouse and wild type mouse brain.

The N-glycoproteins mainly located in the plasma membrane, extracellular region and membrane, while rarely located in the nucleus, cytoplasm, cytoskeleton, and mitochondrion (**Fig. 2G**). The molecular functions of the N-glycoproteins focused on signal transducer activity, transmembrane transporter activity, integrin binding, and receptor binding (**Fig. 2H**). Comparing with the brain proteome, the N-glycoproteins were significantly overrepresented in glutamate receptor signaling, acute phase response signaling, and LXR/RXR activation pathways, *et. al.* (**Figure EV1C**), highlighting the importance of protein N-glycosylation in these pathways. Surface accessibility and secondary structures of the N-glycosites on N-glycoproteins were also depicted (**Figure EV1B**), showing that the N-glycosylation tend to locate at the β-strand more than the α-helix motifs.

### 3. Quantitative N-glycoproteomics revealed depressed protein N-glycosylation in the APP/PS1 mouse model and explored the role of N-glycosylation on NCAM1 membrane localization

In the proteomic analysis, 62 out of 5,046 proteins showed significant changes, of which 47 were up-regulated and 15 were down-regulated in APP/PS1 mice (ratio (AD/WT) ⩾ 1.3 or ⩽ 0.77 and p < 0.05) (**Fig. 3A)**. In this data, 446 proteins were further identified as N-glycosylated proteins **(Fig. 1C)** and none of their protein levels exhibited any difference between wild type and APP/PS1 mice. A total of 38 mouse glycosyltransferases/glycosidases were quantified, while none of them showed any significant changes. The pathways and networks involved in the dysregulated proteins were depicted in **Figure EV2A**. APP, ERK1/2, and EGFR centered network were significantly dysregulated in the APP/PS1 mice. Of the 51 differential N-glycosites, the abundance of 32 N-glycosites increased while that of 19 N-glycosites decreased in APP/PS1 mice (ratio (AD/WT) ⩾ 1.2 or ⩽ 0.83, p < 0.05) **(Figure EV1F)**. Quantitative site-specific N-glycopeptides analysis revealed 289 up-regulated N-glycopeptides on 127 proteins and 729 down-regulated N-glycopeptides on 213 proteins (ratio (AD/WT) ⩾ 1.5 or ⩽ 0.67, p < 0.05) (**Fig. 3B**). In general, the changes in protein expressional level are unremarkable while a sizable portion of site-specific N-glycopeptides were dysregulated and most of them decreased in APP/PS1 mice. These quantification results revealed depressed protein N-glycosylation in APP/PS1 versus wild type mice. The dysregulated N-glycoproteins were significantly enriched in the pathways involving glutamate receptors and other neuronal development and function related pathways (**Fig. 3C**). Many critical membrane proteins, such as glutamate receptors, NCAM1 and 2, Integrins, were heavily glycosylated, and most of their site-specific N-glycans were down-regulated in the AD model (**Fig. 3D**). The glutamate receptor signaling pathway was screened out as most heavily glycosylated pathway (**Figure EV1C**), most of the receptors were N-glycosylated and the majority of their N-glycopeptides were significantly down-regulated in APP/PS1 mice (**Fig. 3E**).

**Figure 3.**
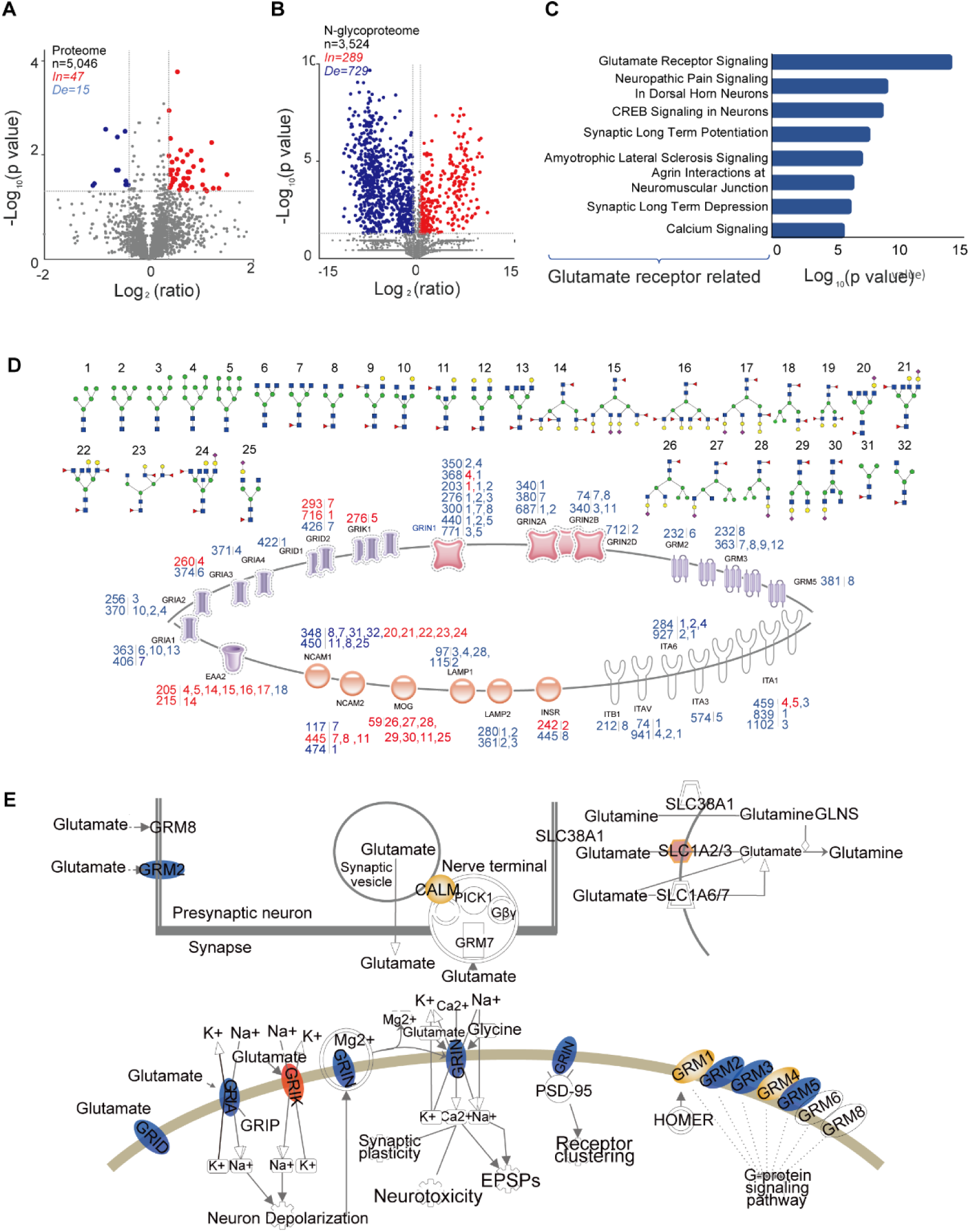
N-glycoproteomic variations of the APP/PS1 mouse versus wild type mouse brain. **(A-B)** Volcano plots showed the differentially expressed proteins and site-specific N-glycopeptides between AD and WT. Blue dots represent the down-regulated proteins or N-glycopeptides while red ones represent the up-regulated proteins or N-glycopeptides. **(C)** The top eight canonical pathways involving the dysregulated N-glycoproteins. **(D)** The identified membrane proteins were depicted. The significantly changed N-glycosites and their N-glycan structures (number 1-32) were shown along with the proteins. Red represents up-regulation and blue represents down-regulation. **(E)** Schematic diagram of the glutamate receptor signaling pathway. The identified proteins were indicated with colored ellipse. Blue ellipses represent proteins with decreased protein N-glycosylation, red ellipses represents increased protein N-glycosylation, and yellow ones represent non-regulated protein N-glycosylation.

To provide a birds-eye view of the multilayered protein N-glycosylations, a circular quantitative N-glycoproteome map was constructed. The quantitative information of 180 proteins with 2132 site-specific N-glycopeptides and 349 N-glycosites that were commonly identified among the proteome, N-glycosite, and site-specific N-glycopeptides analysis were shown (**Fig. 4A)**. The detailed quantitative information of each protein, as well as the glycosites and the site-specific glycopeptides of it can be inquired by zoom-in the corresponding area in the circos plot. The detailed quantitative information of the N-glycosylated metabotropic glutamate receptor 3 (GRM3) were shown as an example. All the identified N-glycopeptides were plotted based on their intensities and the ratios of APP/PS1 versus wild type mice were exhibited (**Fig. 4B**). The abundance of a glycosite was calculated by summing up the intensities of all the glycopeptides on the site. Then the glycosites were plotted according to their intensities and rank of abundances. The up- and down-regulated glycosites were presented in red and blue color, respectively. Most of the down-regulated N-glycosites are in medium to high abundance, including THY1 (site 194), MOG (site 59), glutamate receptors, integrin receptors, and so on (**Fig. 4C**,). To further verify the glycosylation level of identified proteins in N-glycoproteomic analysis, western blots for several representative N-glycoproteins were performed under treatment of PNGase F, Endo H or not (**Fig. 4D and 4E, Supplemental Fig. 3**). Western blot of SHPS1 showed great shift of molecular weight in the PNGase F treated and untreated bands, proving that this protein was heavily glycosylated. The glycosylated (PNGase F-) band in AD sample showed significantly higher intensity than in WT sample, which were consistent with the MS results (**Fig. 4D**). Western blot of THY1 also proved its highly glycosylated status in AD and WT mouse brain, but failed to indicate any variation of glycosylation between the two samples (**Fig. 4E**). This result is understandable since western blot only roughly quantifies the general glycosylation level of protein, while may not represent the glycosylation level at specific sites.

**Figure 4.**
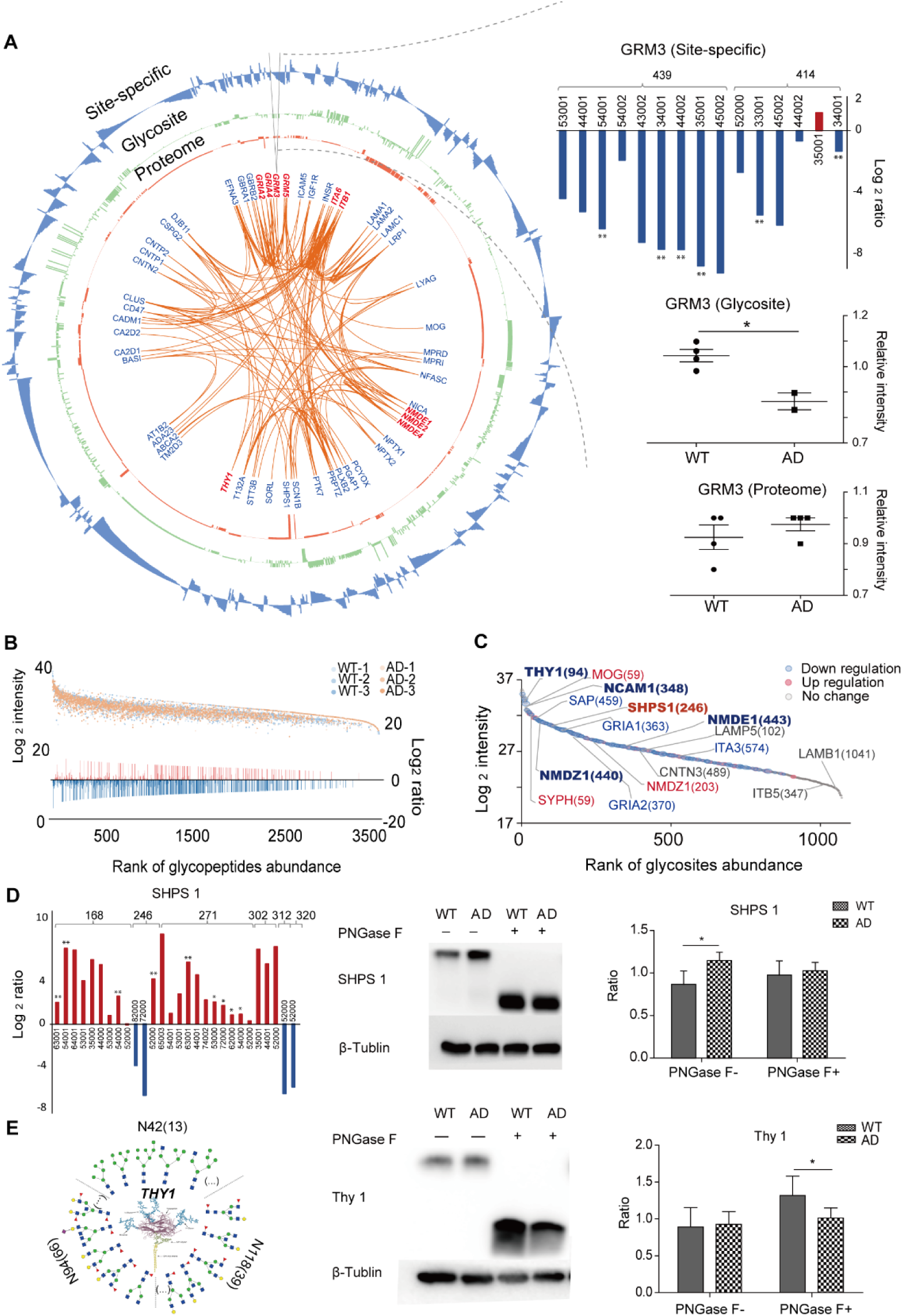
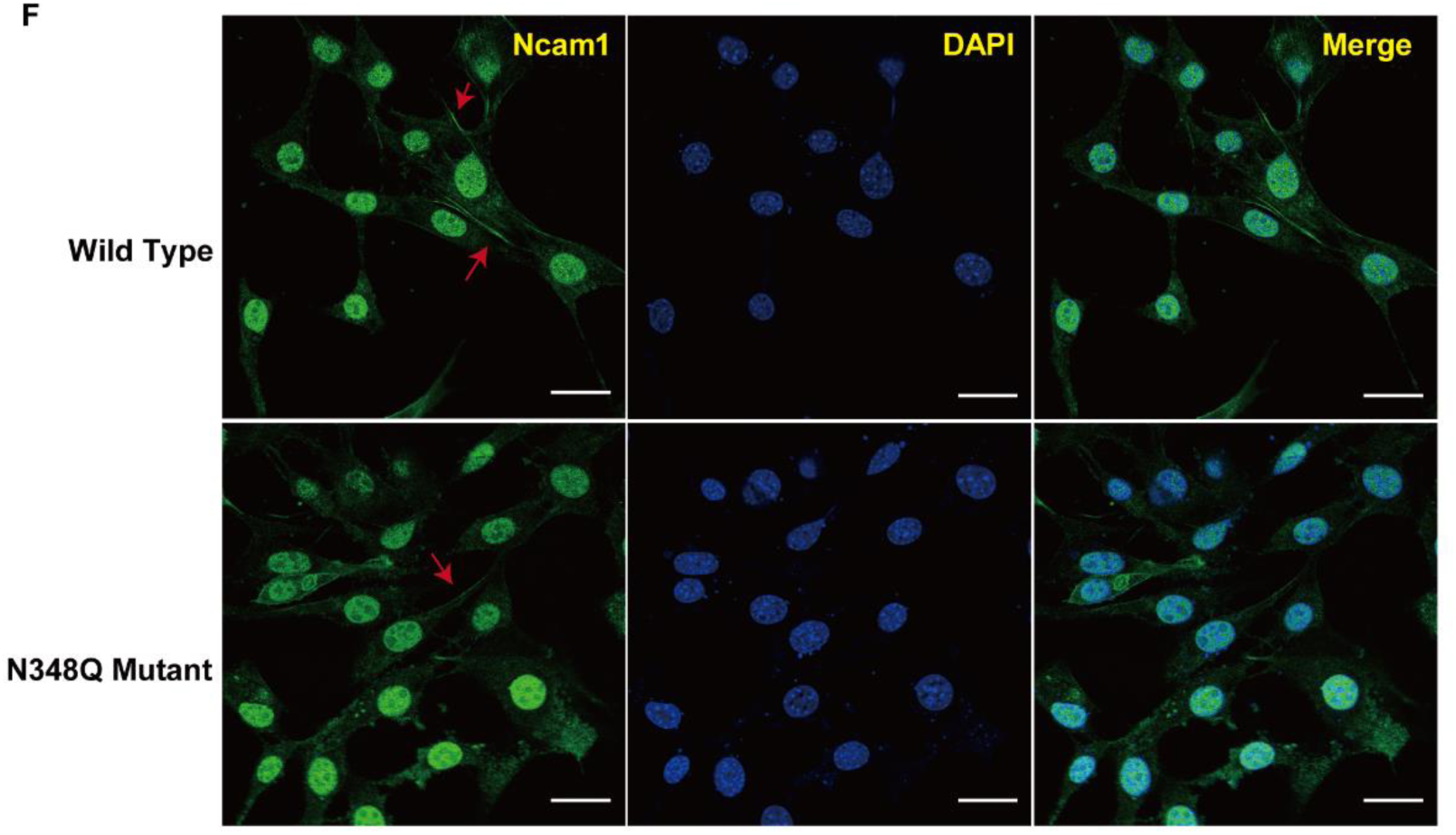
Multilayered quantitative N-glycoproteomes between the APP/PS1 mouse and wild type mouse brain. **(A)** Circos plot visualized the multilayered quantitative information from proteome to N-glycoproteome. The central part of the circos exhibited AD-related proteins with dysregulated N-glycosylation and their interactions with each other. The quantitative results of site-specific N-glycopeptides, glycosite and proteome of GRM3 were showed on the right. Red represents up-regulated N-glycopeptides and blue represents down-regulated ones. The bar results were from triplicate quantitative site-specific N-glycopeptides analysis, quantitative glycosites analysis of quadruplicate of WT and duplicate of AD, or quadruplicate quantitative proteomics analysis. **(B)** All the identified N-glycopeptides were plotted based on their intensities and the ratios of WT versus AD. Ranking of the N-glycosites were calculated by summing the intensities of their corresponding site-specific N-glycopeptides from AD and WT mice. **(C)** Some of the brain proteins with dysregulated N-glycosites are marked. The size of the bubble is inversely proportional to p values. Red represents up-regulated N-glycopeptides and blue represents down-regulated ones. **(D)** The quantitative results of site-specific N-glycopeptides on six sites of SHPS1 were showed. Red represents up-regulated N-glycopeptides and blue represents down-regulated ones. Validation of protein glycosylation were achieved by western blots. The bar results were from triplicate quantitative analysis using Quality One. **(E)** Example of a hyperglycosylated protein. Multiple N-glycans were attached on each site of Thy1_MOUSE. Validation of protein glycosylation were achieved by western blots. The bar results were from triplicate quantitative analysis. **(F)** The exogenous NCAM1 (wild type) or NCAM1 construct (N348Q mutant) was transfected into HT22 cells for 48h. Wild type and N348Q mutation samples were immunostained with anti-Ncam1 (green). Nuclei was visualized with DAPI (blue). Ncam1 mostly located in cell membrane and this was disrupted by the N348Q mutation in Ncam1. Up panels show wild type, and down panels show N348Q mutant. Scale bars, 20 μm. All data shown represent mean ± SEM. * The equivalent of p <0.05 and ** the equivalent of p <0.01; Student’s t-test was used for the data analysis.

Neural cell adhesion molecule (NCAM), also called CD56, is a homophilic binding glycoprotein expressed on the surface of neurons, glia and skeletal muscle. NCAM1 and NCAM2 are two paralogs of NCAM. NCAM1 is a cell adhesion molecule involved in neuron-neuron adhesion. During brain development, NCAM1 is heavily polysialylated, which determines its adherent properties. The polysialylated modification of NCAM1 modulate inputs of brain circuitry or activity of ion channels(Mehrabian *et al*, 2016; Mirnics *et al*, 2005). In this work, we identified multiple glycopeptides on NCAM1 and NCAM2, including some sialylated ones (**Fig. 3D**). Site 348 of NCAM1 was one of the most heavily glycosylated site, and its glycosylations were down regulated in the APP/PS1 sample **(Fig. 4D**). To investigate the function of NCAM1, we overexpressed the exogenous NCAM1 (WT) or glycosylation-site mutant in the mouse hippocampal neuronal cell line HT22. The mutation was generated by replacing the asparagine residue at site 348 with glutamine (N348Q) of NCAM1. As shown in confocal images, WT NCAM1 expressed membranous and cytoplasmic locations. However, overexpressing N348Q mutant of NCAM1 tended to locate in cytosol rather than membrane in HT22 cells (**Fig 4F**). This result showed that the glycosylation of the Asn^348^ were crucial for the membrane localization of NCAM1. And depressed Asn^348^ glycosylation, as occurred in the APP/PS1 sample, would cause the loss of function of NCAM1 as a membrane protein.

### 4. Down-regulated glycans and glycan patterns in the APP/PS1 sample and the effects of blocking the surface glycans on hippocampal neurons physiology

The glycomes of the APP/PS1 and wild type mouse brain were acquired by two strategies. Lectin microarray containing 91 lectins successfully recognized 83 different glycan epitopes on brain proteins in APP/PS1 and wild type mice (**Fig. 5A, B and Figure EV4A, B**). About 70 kinds of lectins showed decreased trends and 13 lectins were significantly down-regulated in APP/PS1 mice (ratio≥1.2 or ≤ 0.83, p<0.05) (**Fig. 5A, B, and Table 1 and Figure EV4A**). The signals for AAL, PSA and LBA were significantly decreased in the APP/PS1 mice, indicating the decline in fucosylated glycans in APP/PS1 mice which is reported for the first time. The signal for WGA binding GlcNAc and multivalent sialic acid residues decreased as well, which agreed with previous studies (Fodero *et al*, 2001; Saez-Valero *et al*, 1999). Other lectins with altered binding intensities included the mannose, galactose, GalNAc, and complex type binding lectins **(Fig. 5B)**. We also independently blotted 14 lectins in APP/PS1 and wild type mouse brain. The staining intensities of AAL, WGA, Jaclin, and PSA were significantly lower in APP/PS1 mice (**Fig. 5C)**. Another 10 lectins, such as PHA-L, DSL, and SBA, showed no significant changes **(Figure EV4E)**. These lectin blot results are highly consistent with those from the microarray data.

**Figure 5.**
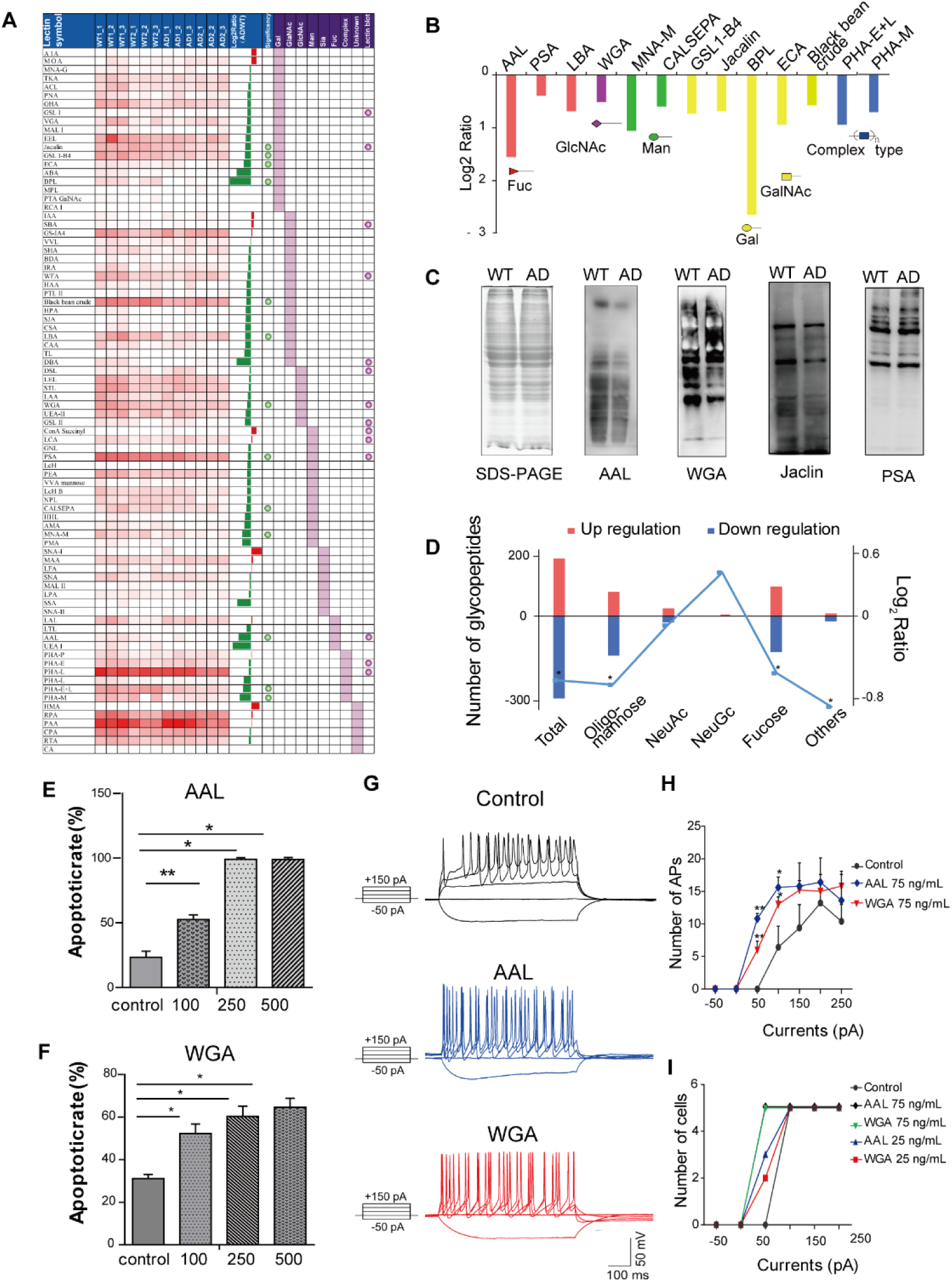
AAL and WGA Treatment induced increased apoptosis and hyperexcitability of hippocampal neurons. **(A)** An overview of the results of the lectin microarray. Two biological replicates and three technical replicates were performed from both AD and WT mouse brain proteins. The net intensities of each lectin from the six replicates were shown with a color gradient. The log2-transformed ratios of AD/WT were sorted in a descending order. The significantly changes lectins were indicated with a ball. Green ball means the decreased lectin binding. All of the lectins in lectin microarray were classified into seven categories based on the main monosaccharides binding specificity. The lectins which were validated by lectin blots were indicated with purple balls. **(B)** The thirteen lectins with significantly decreased signals in AD. The lectins were arranged on the basis of sugar-binding specificities of lectins as follows: Fuc, fucose binder; GlcNAc, α- or β-GlcNAc binder; Man, mannose binder; Gal, α- or β-Gal binder; GalNAc, α- or β-GalNAc binder. **(C)** The lectin blotting analysis of AAL, WGA, Jaclin and PSA. The staining intensities of AAL, WGA and Jaclin were lower in AD mice than that in WT mice while the staining intensities of PSA did not alter in AD mice. The same amount of brain proteins (20 μg) from AD and WT mice were loaded onto 8% SDS-PAGE for lectin blotting. **(D)** The numbers of the significantly changed site-specific N-glycopeptides with different N-glycan subcategories were showed as histogram. The total intensities of N-glycopeptides with different N-glycans were summed and compared in AD versus WT, and plotted as line chart. **(E-F)** Apoptosis of hippocampal neurons after treatment with PBS, 100 ng/mL, 250 ng/mL, and 500 ng/mL AAL or WGA were assessed using PI staining. The apoptosis rate of AAL groups (E) and WGA (F) were quantitated (2 or 3 samples per treatment group and 3 views per sample were counted). **(G)** Representative traces of induced action potentials (AP_s_) from −50 pA to 150 pA injections for normal hippocampal neurons of rat as control (blank), treated with 75 ng/mL AAL for 24 h as AAL group (blue) and 75 ng/mL WGA for 24 h as WGA group (red). **(H)** Statistical analysis of AP frequency in all three groups. **(I)** The AP firing threshold of hippocampal neurons in five groups with different lectin concentration. n=5 neurons for each individual. All data shown represent mean ± SEM. * The equivalent of p <0.05 and ** the equivalent of p <0.01; Student’s t-test was used for the data analysis.

The oligosaccharide compositions of protein-attached N-glycans were also identified and quantified in the site-specific N-glycoproteomic analysis, and 290 kinds of N-glycans were identified (**Fig. 5C)**. The abundance levels of the N-glycans were analyzed based on the summed intensities of their corresponding site-specific N-glycopeptides. The abundances of the total, oligo-mannose, fucosylated N-glycans and sialylated N-glycans (Neu5Ac and Neu5Gc) were analyzed, and the total, oligo-mannose, fucosylated N-glycans showed significantly decrease trends in the APP/PS1 mice (**Fig. 5D)**. In addition, the overall lengths of N-glycans in APP/PS1 mice were found to be a little longer than that in wild type mice (**Figure EV4B)**.

Cellular component analysis showed that fucose, oligo-mannose, NeuAc and NeuGc containing proteins were mainly located in the plasma membrane and extracellular region (**Figure EV4C**). The above results, together with the lectin microarray and lectin blot data, demonstrated the significant depressed N-glycome in APP/PS1 mice from different angel, especially for the fucosylated and oligo-mannose glycoforms.

To further examine whether the protein glycosylation impairment was associated with neuron’s pathological development in the APP/PS1 moues model, the reduction of protein glycosylation on neuron surface was mimicked by incubating hippocampal neuron cells with AAL and WGA lectins, which can block fucosylated glycans and GlcNAc residues, respectively and were both down-regulated in the lectin array analysis. Rat hippocampal neurons were incubated with PBS (control), different doses of AAL or WGA lectins for 24 h followed by propidium iodide (PI) staining. The apoptosis rate of hippocampal neurons increased significantly from 31.1% to 52.2% with 100 ng/mL and to 70% with 500 ng/mL AAL treatment (p<0.05) (**Fig. 5E, and Figure EV5A)**. The results indicated that blocking fucosylation with AAL severely suppressed the viability of hippocampal neurons. Similar results were observed in the WGA treatment experiments, where the apoptosis rate significantly increased from 23.3% to 52.5% with 100 ng/mL and up to 98.9% with 500 ng/mL WGA treatment (p<0.01) (**Fig. 5F, and Figure EV5B**).

Electrophysiological analysis was then conducted in lower doses of AAL and WGA that can maintain neuron viability. Hippocampal neurons were treated with PBS, AAL or WGA of 25 ng/mL or 75 ng/mL for 24 h. The current clamp recordings showed that AAL and WGA treated cells exhibited higher action potential (AP) frequency in response to +100 pA and +150 pA current injections compared with the control (**Fig. 5G**). The treated neurons reached AP threshold under lower current injections (+50 pA) than the control (+100 pA). The number of excitatory neurons at +50 pA current injection increased with increasing lectin concentration, exhibiting a dose dependency (**Fig. 5H**). These results demonstrated that blocking neuron surface glycans with AAL and WGA induced the hyperexcitability of hippocampal neurons. It has been reported that neuronal hyperactivity is an early phenotype in Alzheimer’s disease and could trigger apoptosis of neurons(Abiega *et al*, 2016; Nuriel *et al*, 2017). Our results demonstrated that declined glycosylation on the cell surface of hippocampal neurons can induce hyperexcitability and further apoptosis, which implied the important role of glycosylation in the pathogenesis of the Alzheimer’s disease.

## DISCUSSION

There are urgent needs for global and site-specific analysis of glycoproteins in complex biological samples for understanding of glycoprotein functions and cellular activities. However, it is extraordinarily challenging because of the low abundance of many glycoproteins and the heterogeneity of glycan structures. In this study, we applied the pGlyco 2.0 strategy to profile the N-glycopeptides in APP/PS1 and wild type mouse brains and successfully identified 3,524 site-specific glycopeptides. This data provided comprehensive characteristics of the N-glycoproteins of mouse brain and fully demonstrated the complexity and heterogeneity of N-glycosylation modifications (**Fig. 2**).

To further elucidate the protein glycosylation changes in APP/PS1 versus the wild type mice, we combined data from the proteome, N-glycosites, and glycome with that from intact N-glycopeptides and constructed the multilayered N-glycoproteome profiles. The proteome changes in the brain of APP/PS1mice were relatively small, while the N-glycoproteome profiling showed significant down-regulations in APP/PS1 mice. The N-glycoproteome, glycome and lectin blotting experiments demonstrated the same down-regulation trends of protein glycosylation in the APP/PS1 mouse model from different angles. Complicated site-specific variations were observed in some glycoproteins and their sites, such as SYPH (site 59) and AT1B1 (site 158) shown in **Figure EV2G.** These two sites both have multiple N-glycans attached with different abundance and diverse changing trends. It’s not clear whether these site-specific N-glycans with different trends have independent or redundant functions.

Most of the membrane proteins, such as glutamate receptors, were found to be highly N-glycosylated and their glycosylation levels are consistently down-regulated in APP/PS1 mice. L-glutamate is the major excitatory transmitter in vertebrate central nervous system (CNS). The excitatory responses of glutamate are mediated by a number of functionally distinct cell-membrane receptors and almost every glutamate receptor subtype has been implicated in mediating excitotoxic neuronal cell death(Sattler and Tymianski, 2001). N-glycans have been reported to affect maximal currents and desensitization of AMPA-type glutamate receptors and kainite receptors and are required for folding or trafficking of NMDA receptors(Panin, 2014). This study offered rich and detailed N-glycosylation of 14 glutamate receptors and revealed the depressed N-glycosylation of glutamate receptors in APP/PS1 mice (**Fig. 3C, D, and E**). The dysregulated N-glycosylation of these glutamate receptors and other membrane proteins may interfere their normal functions and play roles in Alzheimer’s disease pathological process. From the perspective of the glycome analysis, the overall N-glycans, oligo-mannose N-glycans, as well as the fucosylated N-glycans were all significantly down-regulated in the APP/PS1 mice. Lectin microarray and lectin blotting experiments revealed decreased signals in the corresponding lectin signals such as the AAL recognizing fucose and WGA recognizing GlcNAc. Our data presents the overall down-regulation trends and the detailed site-specific changes of the N-glycosylation modifications in APP/PS1 mice. Considering none of the 38 identified glycosyltransferases/glycosidases were dysregulated in our data, we have no idea of the causal factors for the down-regulated glycosylation. We speculated that dysregulated enzymes, and metabolic disorders of glucose and other monosaccharides are both possible factors inducing the depressed glycosylation.

To explore the influence of the impaired glycosylation on protein functions and cellular activities, we generated exogenous NCAM1, including WT NCAM1 and the N348Q mutant in HT22 cells and blocked the surface glycans of rat primary hippocampal neurons with AAL and WGA lectins. In accordance with our expectations, replacing the Asn^348^ to glutamine and removing the glycans on this site impaired the membrane localization of NCAM1. We believe that the effects of N-glycosylation on the localization and functions of related proteins are ubiquitous. And generally depressed N-glycosylation would cause a systematic disorder of these proteins. Lectin incubation showed that both AAL and WGA can cause excitability and activation of neuronal cells and further led to neuronal apoptosis. This result demonstrated the important role of the overall protein N-glycosylation in maintenance of the functions and survival of neuronal cells. Based on the glycoproteomic findings, we created a schematic model summarizing the possible causes of the depressed N-glycosylation and we think the depressed N-glycosylation may affect the localization, function, and stability of a large number of glycoproteins, and may even lead to cytotoxicity of the neuron (**Fig 6**). These consequences would further exacerbate the AD pathology. Currently, we are performing experiments and trying to explore more influences of N-glycosylation on proteins and neuron cells.

**Figure 6.**
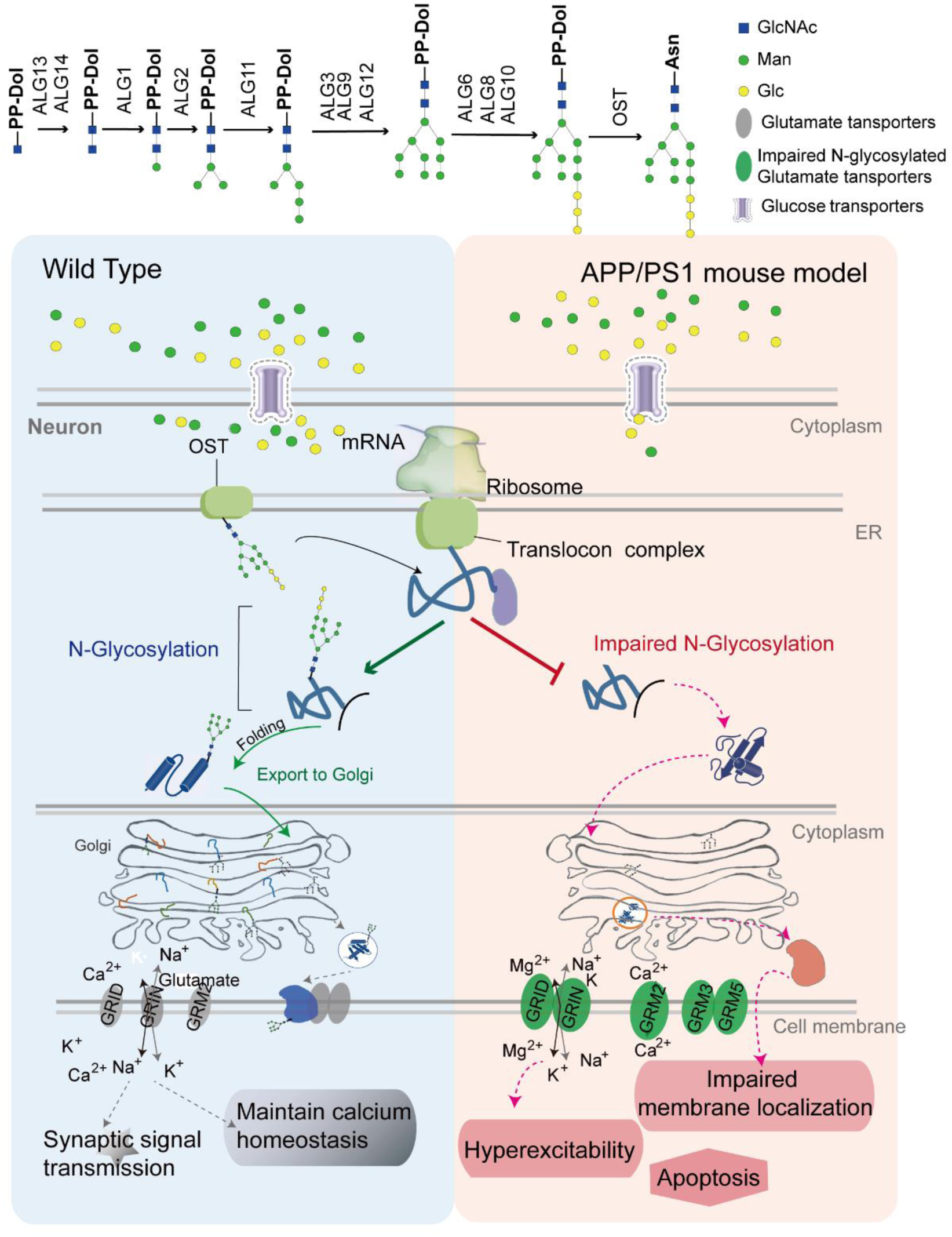
Schematic model. Schematic model summarizing the findings of the multilayer N-glycoproteomic profiles. The pathways and enzymes involved in glycosylation were shown in the upper panel. Under normal conditions, proteins are glycosylated and correctly folded in the endoplasmic reticulum and maintain normal physiological functions, such as synaptic signal transmission and calcium homeostasis; however, protein N-glycosylation is impaired, and important proteins such as glutamate receptor GRM3 or cell adhesion molecule NCAM1 cannot work normally in APP/PS1 mice (AD model), which resulted in hippocampal neurons membrane localization damage, hyperexcitability and apoptosis.

Taken together, this study revealed the generally impaired protein N-glycosylation in APP/PS1 mice through a comprehensive profiling of the multilayered N-glycoproteome, and further demonstrated the potential roles of the depressed N-glycosylation on glycoproteins and neuron cells in Alzheimer’s disease.

## MATERIALS AND METHODS

### 1. Alzheimer’s disease model

Brain tissues from the APP/PS-1 double-transgenic mice and wild type mice, aged 12 months, were kindly presented by Professor Chunjiu Zhong from Zhongshan Hospital.

### 2. Lectin microarray

The lectin microarray was manufactured as previously described (Tao *et al*, 2008) and provided by Wayen Biotechnologies (Shanghai, China).

### 3. Glycopeptide enrichment

Labeled or unlabeled peptides were enriched by ZIC-HILIC method.

### 4. Liquid chromatography-mass spectrometry (LC-MS) for iTRAQ-labeled samples

Peptides labeled by iTRAQ were pre-fractionated with high pH reverse phase LC. Each fraction was analyzed on a Triple-TOF^TM^ 4600 system (AB SCIEX, USA) equipped with a nano-HPLC (Eksigent Technologies). The iTRAQ-labeled de-glycopeptides were analyzed on an Orbitrap Fusion Tribrid system (Thermo Fisher Scientific, USA).

### 5. Database searching for iTRAQ-labeled peptides and de-glycopeptides

The iTRAQ-labeled peptides were identified with Proteome Discoverer (1.4.0.288) using Mascot against the SwissProt mouse database (2015_03, mouse, 16,711 entries).

### 6. LC-MS analysis of site-specific N-glycopeptides

Enriched glycopeptides were directly analyzed by nano spray LC-MS/MS on an Orbitrap Fusion Tribrid (Thermo Scientific) without the trap column. The gradient was 4 hours in total for complex samples. Precursors were fragmented under a stepped collision energy.

### 7. Database searching for the analysis of site-specific N-glycopeptides

Site-specific glycopeptides were interpreted using the software pGlyco 2.0. The glycan database was extracted from GlycomeDB (www.glycome-db.org). Trypsin and protein databases with species of Mus musculus (16,711 entries) were used. Quantification of site-specific glycopeptides were based on the peak areas as previously described (Schwanhausser *et al*, 2011).

### 8. Hippocampal Neurons Culture

Cultures of hippocampal neurons were prepared from P0 Sprague–Dawley (SD) rat as previously described (Dotti *et al*, 1988). Procedures involving animals and their care were performed in accordance with the Animal Care and Use Committee of the Institute of Neuroscience.

### 9. Cell viability and electrophysiological recordings

The hippocampal neurons of rat were treated with different concentrations of AAL and WGA lectin treatment for 24 hours on the 7th day after attachment.

### 10. Statistical analysis

A two-tailed student’s t-test was used for significance estimation in all quantification experiments, and *P* values less than 0.05 were considered significant.

### 11. Data availability

The proteome, N-glycoproteome, and glycome data is accessible in http://firmiana.org^34^. Acquired data are available from the corresponding author upon reasonable request.

## ACKNOWLEDGMENTS

Thanks to Professor Xuewen Cheng from SIBS (Institute of Neuroscience, Shanghai Institutes for Biological Sciences, Chinese Academy of Sciences) for his advice and help on the functional verification of hippocampal neurons.

## AUTHOR CONTRIBUTIONS

P.F. designed and performed the N-glycoproteomic experiments, data analysis and prepared the manuscript. J.-J.X designed and performed the biological validation experiments, data analysis and prepared the manuscript. S.-M.S provided the animal models and contributed to the data analysis. M.-Q.L. performed the data analysis and contributed to the manuscript preparation. L.-J.Y. contributed to the experiments. Y.-T.X. performed the data analysis. L.Z., X.G., G.-Q.Y, and J.Y. performed the MS/MS analysis. W.-J. Q. and Z.-F. W. provided the kindly help of electrophysiological recordings experiments. Y.Z. performed the data analysis. P.-Y.Y. directed the experiments and revised the manuscript, and H.-L.S. directed the project and revised the manuscript.

## COMPETING INTEREST

The authors report no competing interests.

## FUNDING

This study was supported by the National Key Research and Development Program of China (2018YFA0507501, 2017YFA0505001), Shanghai Science and Technology Research Project (16ZR1402400), and the Zhuo Xue Program of Fudan University.

## SUPPLEMENTARY MATERIAL

Supplementary material is available online.

## EXPANDED VIEW FIGURE LEGENDS

**Figure EV1.**
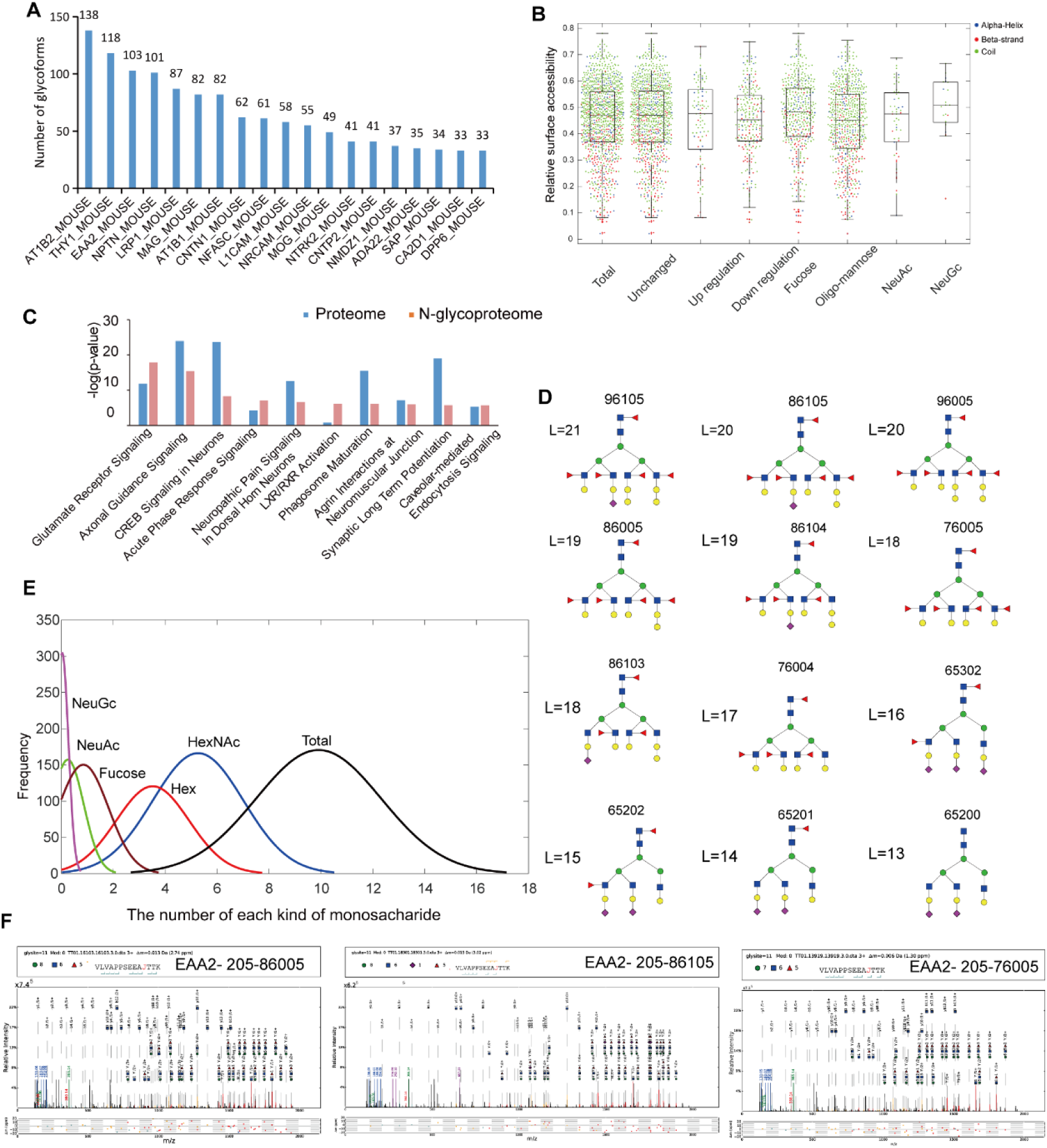
Characteristics of the proteome, N-glycoproteome, and glycome in mouse brains. **(A)** The top 20 proteins with the highest number of site-specific N-glycopeptides on them. **(B)** The relative accessibility of N-glycosites on the alpha helix, beta strand, and coil. **(C)** Pathway comparison of the proteome and N-glycoproteome using IPA. **(D)** Speculated structures of long N-glycans identified in this work with 13-21 monosaccharides. **(E)** The distribution of each kind of monosaccharide in site-specific glycoproteome. **(F)** Examples of the MS/MS spectra of the N-glycopeptides on site 205 of EAA2_MOUSE with “86005”, “86105” and “76005” N-glycans.

**Figure EV2.**
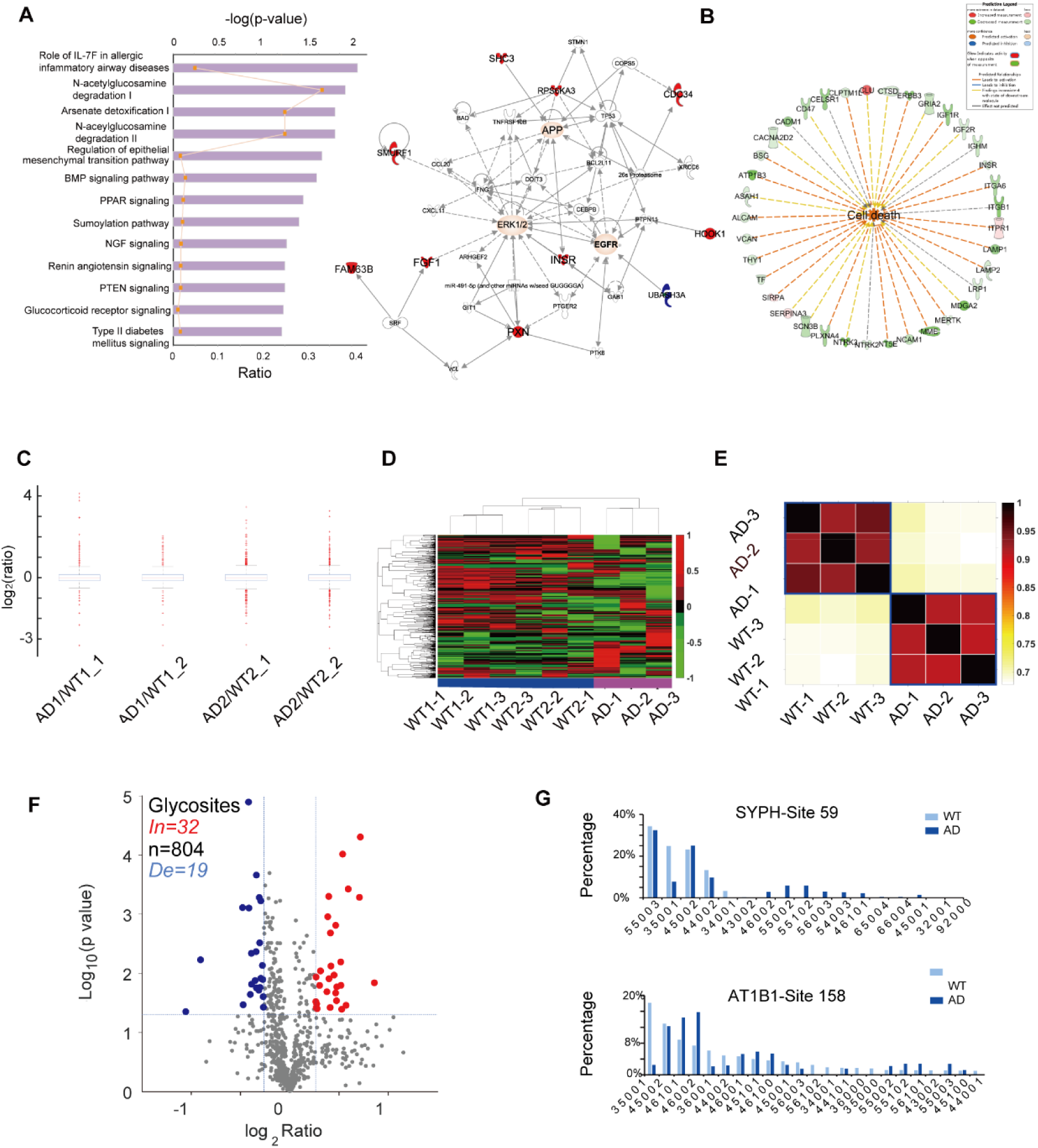
Quantitative analysis of the multilayered N-glycoproteome in AD and WT mouse brain. **(A)** The enriched biological pathways and molecular networks of differentially expressed proteins in AD mice. Red plots represent up-regulated proteins while blue ones represent down-regulated proteins. **(B)** Green indicates down-regulated, red, up-regulated. The color intensity is correlated with fold change. Straight lines are for direct gene to gene interactions, dashed lines are for indirect ones. **(C)** Boxplot indicates the distribution of ratios of all quantified proteins in global proteome. Two biological replicates and two technical replicates were performed. **(D)** Cluster analysis of N-glycosites quantification. **(E)** Pearson correlation coefficients depicted the intensities of site-specific N-glycopeptides from triplicates experiments. **(F)** Volcano plots showed the differentially expressed N-glycosites. Blue dots represent the down-regulated N-glycosites while red ones represent the up-regulated N-glycosites. **(G)** The site occupancy of different N-glycans on the site 59 from SYPH and the site 158 from AT1B1 in AD and WT were exhibited.

**Figure EV3.**
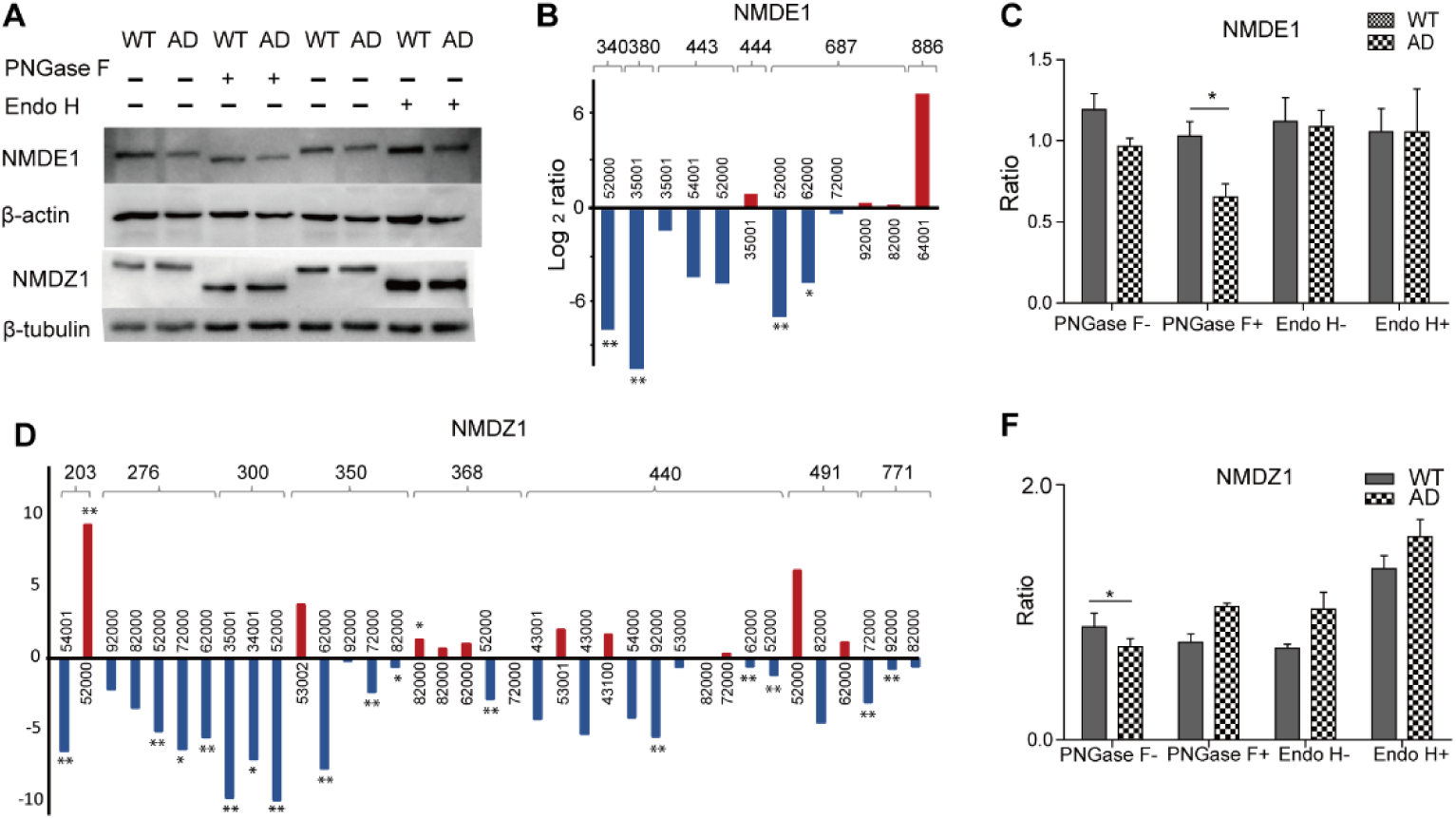
Validation of protein glycosylation using western blots. The proteins from AD and WT mouse brain were treated with or without PNGase F or Endo H. **(A)** Proteins were separated on SDS-PAGE and analyzed by western blots. Note that NMDE1 and NMDZ1 displayed reduced molecular weight compared with glycosylated proteins. **(B-C)** The quantitative results of site-specific N-glycopeptides on six sites of NMDE1 were showed. **(D-F)** The quantitative results of site-specific N-glycopeptides on six sites of NMDZ1 were showed. Red represents up-regulated N-glycopeptides and blue represents down-regulated ones. The bar results were from triplicate quantitative site-specific N-glycopeptides analysis. means p <0.05 and ** means p <0.01.

**Figure EV4.**
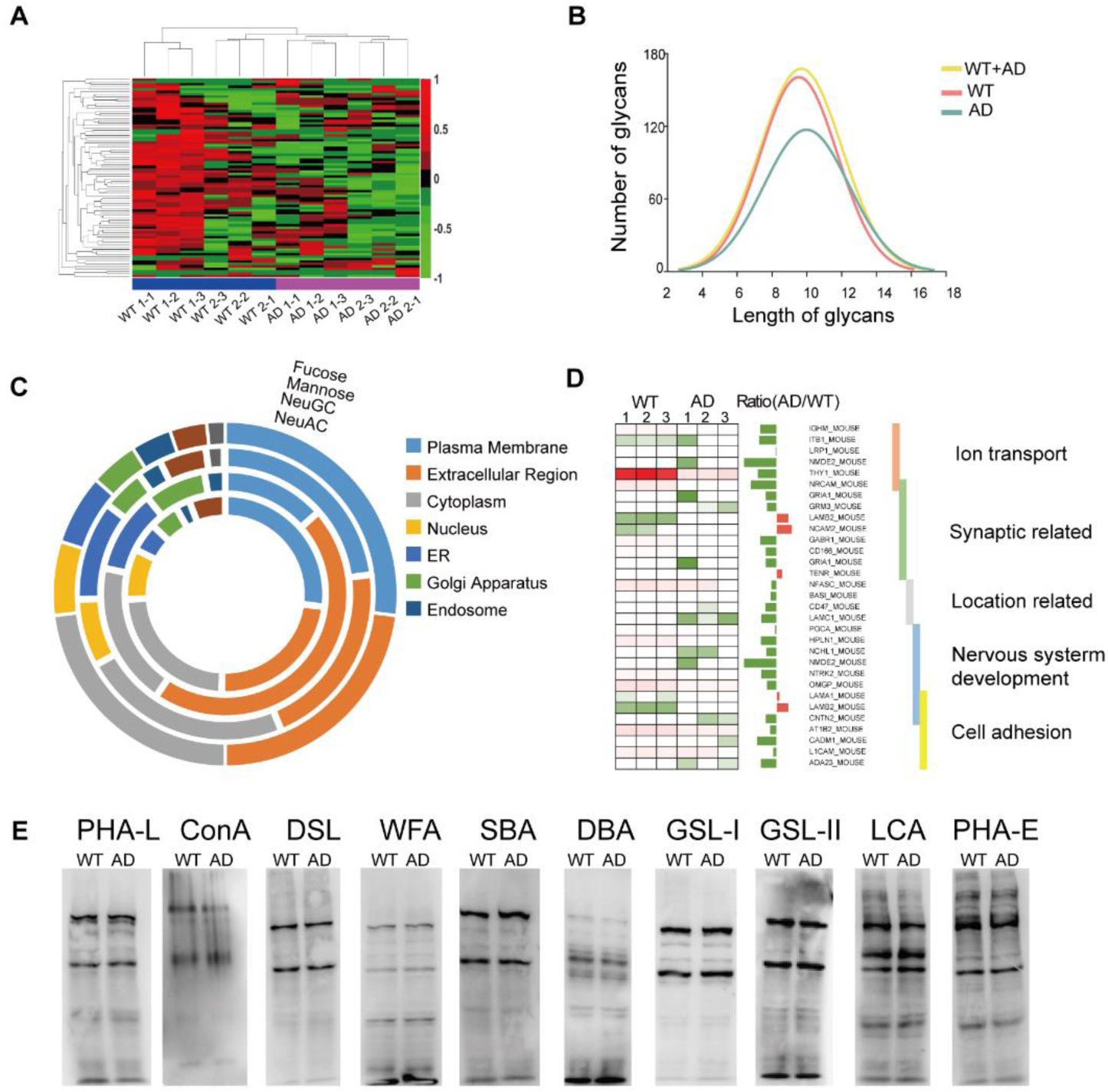
Glycan profiling of mouse brain proteins indicated the differential glycan patterns between AD and WT mice. **(A)** Heat map analysis of lectins with positive binding. The samples are listed in columns, and the lectins are listed in rows. **(B)** The distribution of lengths of N-glycans in AD and WT. **(C)** The cellular localization distribution of glycoproteins containing fucose, mannose, Neu5Ac or Neu5Gc. **(D)** Molecular function of the glycoproteins with down-regulated fucosylated site-specific N-glycopeptides. **(E)** Results of lectin blotting analysis. The staining intensities of PSA and Jacalin were lower in AD mice than that in WT mice, while the staining intensities of the other lectins did not alter in AD mice. The same amount of brain proteins (20 µg) from AD and WT mice was loaded onto 8% SDS-PAGE for lectin blotting.

**Figure EV5.**
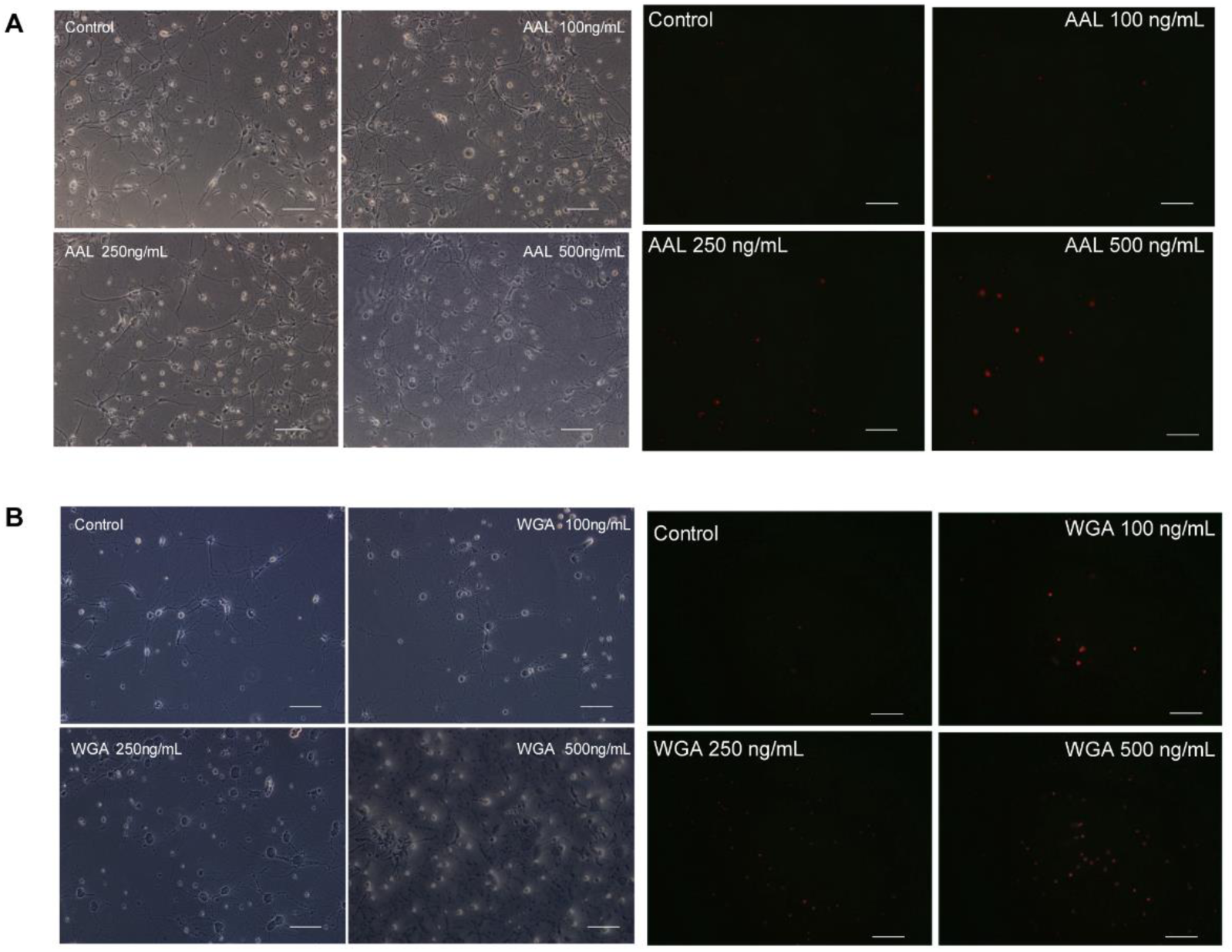
Treatment with AAL and WGA induced increased apoptosis of hippocampal neurons. **(A-B)** Microscopy and fluorescence images of rat hippocampal neurons after treatment with different concentrations of AAL (a) or WGA (b)for 24 hours on the 7th day after attachment. Length of scale bar equals 50 µm.

